# Multiscale modeling of tumor adaption and invasion following anti-angiogenic therapy

**DOI:** 10.1101/2021.08.25.457537

**Authors:** Colin G. Cess, Stacey D. Finley

## Abstract

In order to promote continued growth, a tumor must recruit new blood vessels, a process known as tumor angiogenesis. Many therapies have been tested that aim to inhibit tumor angiogenesis, thus starving the tumor of nutrients and preventing tumor growth. However, many of these therapies have been unsuccessful and can paradoxically further tumor development by leading to increased local tumor invasion and metastasis. In this study, we use agent-based modeling to examine how hypoxic and acidic conditions following anti-angiogenic therapy can influence tumor development. Under these conditions, we find that cancer cells experience a phenotypic shift to a state of higher survival and invasive capability, spreading further away from the tumor into surrounding tissue. Although anti-angiogenic therapy alone promotes tumor cell adaptation and invasiveness, we find that augmenting chemotherapy with anti-angiogenic therapy improves chemotherapeutic response and delays the time it takes for the tumor to regrow. Overall, we use computational modeling to explain the behavior of tumor cells in response to anti-angiogenic treatment in the dynamic tumor microenvironment.

## 1 INTRODUCTION

It has long been known that angiogenesis, the formation of new blood vessels, is a feature common to almost every solid tumor. As the tumor grows, it depletes its local environment of oxygen (called “hypoxia”), and the subsequent response to hypoxia is the secretion of various pro-angiogenic factors, such as vascular endothelial growth factor (VEGF). These factors recruit blood vessels to the tumor, resupplying nutrients and removing waste products. It has also been found that angiogenesis promotes tumor metastasis to distal sites [1,2]. Therefore, being able to modulate tumor angiogenesis is seen as a way to potentially provide some therapeutic benefit. Since the discovery of tumor angiogenesis, various therapies have been developed that primarily target the effects of pro-angiogenic factors, with the aim of preventing vascular recruitment [3,4]. These therapies work by starving the tumor of nutrients, preventing continued growth and killing the tumor. However, the therapies provide limited actual clinical benefit [5,6]. In addition, clinical and mouse studies have found that inhibiting angiogenesis can not only fail to remove the tumor, but also lead to a more invasive and metastatic tumor [7–9].

To understand why invasion and metastasis occur following anti-angiogenic therapy, we must look at how the tumor responds to the harsh microenvironmental conditions (primarily hypoxia and lowered pH) that occur when its blood supply is cut off. Hypoxia has been shown to lead to other adaptive changes in cancer cells besides angiogenic factor secretion, notably an upregulation of glucose transporters, which contributes to the Warburg Effect [10–13] and ultimately leads to elevated glucose metabolism and increased production of lactic acid as a byproduct. This lactic acid secretion helps the cancer cells invade into nearby tissue by killing normal cells and degrading extracellular matrix [14,15]. Hypoxia has also been found to promote cancer cell migration and epithelial-mesenchymal transition [16,17], further increasing the cells’ invasive capabilities. In addition to hypoxia-mediated effects, cancer cells adapt to the acidic conditions produced by increased glycolysis, allowing them to survive at a pH that would kill normal cells [14,18]. These adaptations, which are a result of lowered blood supply, explain the *in vivo* and clinical observation that tumor invasion is advanced by anti-angiogenic treatment.

In order to explore the tumor in a more detailed setting, computational models have been used to simulate tumor angiogenesis. These models allow for many different perturbations to be tested and a vast amount of information examined. These studies would otherwise be very prohibitive to perform *in vitro* or *in vivo*. While many different types of models have been developed to examine angiogenesis, ranging from intra-cellular signaling models to whole tumor models [19–24], we will focus here solely on a type of model known as the agent-based model (ABM). ABMs model cells as discrete individuals, capturing complex cell-to-cell interactions and spatial morphologies. To date, a plethora of ABMs have been developed to examine various aspects of tumor vasculature [25–32]. These models focus on a variety of different aspects of angiogenesis, vascular properties, and microenvironmental conditions. Depending on the model focus, vasculature has been treated as simply as point sources or as complexly as explicit vessel structure with detailed blood rheology. Together, these models have advanced understanding of tumor-vasculature dynamics by allowing researchers to explore in great detail how modulating different interactions leads to differences in the temporal and spatial development of the tumor ecosystem.

In this study, we aim to use agent-based modeling to better understand the mechanisms that contribute to more invasive and metastatic tumors following anti-angiogenic treatment. While previous models on this subject focus mostly on the interplay between tumor and vasculature, they often neglect how cancer cells change with time in response to their dynamic environment, which is the focus of our work here. Using a simple approach to simulate the vasculature within an ABM, we focus on how tumor cells adapt to harsh microenvironmental conditions and how reducing angiogenesis leads to changes in tumor phenotype and spatial morphology. In addition, we focus on an effect of anti-angiogenic treatment known as vascular normalization and how it augments the effects of chemotherapy by transiently increasing blood flow. Overall, we find that anti-angiogenic therapy leads to tumor adaptation and a phenotypic shift towards more invasive properties: increased glycolysis, increased resistance to acidic pH, and increased migration, leading to invasion of the surrounding tissue. However, we also find that augmenting chemotherapy with anti-angiogenic therapy improves the chemotherapy’s ability to suppress tumor growth.

## 2 METHODS

### Model overview

Our model is an agent-based model representing tumor growth in a 2D tissue slice. It contains two main entities, cancer cells and blood vessels, along with several diffusible factors. An overview of the model is displayed in Figure 1. The model is constructed in C++ and can be found at: https://github.com/FinleyLabUSC/Anti-angiogenic-treatment-ABM-model.

**Figure 1.**
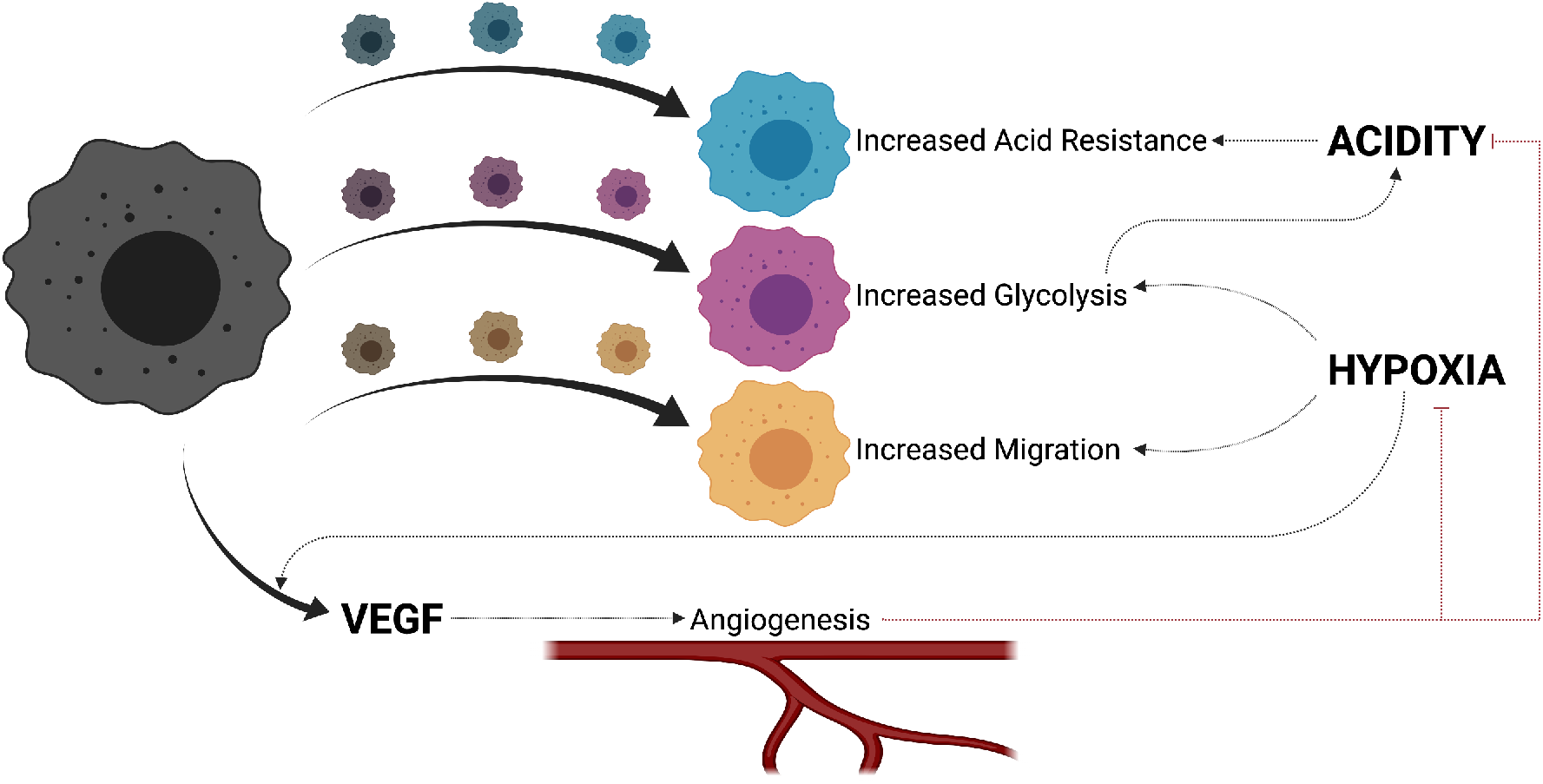
Model schematic. Cancer cells start at the base phenotype (black) and adapt phenotypically to environmental conditions. Acidity causes an increase in acid resistance (blue), while hypoxia causes increases in glycolysis (purple) and migration (yellow). Hypoxia also causes VEGF secretion by cancer cells, which promotes angiogenesis and limits hypoxia and acidity.

### Cellular forces

#### Inter-cellular forces

Cells are modeled individually and are represented by a point, *x*_*i*_(*t*), and a radius. Adjacent cells exert forces on each other, based on the assumption that cells are deformable and can interact with each other. Following the method presented by Osborne et al. [33], we use Equation 1 to model the effects of forces on cells, numerically solving it via Equation 2.

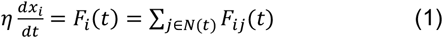

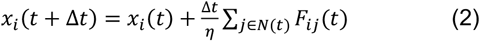

In these equations, *F*_*ij*_(*t*) is the force exerted on cell *i* by cell *j, N*(*t*) is the number of cells in the environment, and *η* is the damping coefficient.

There are two types of forces between cells: repulsive, if the cells are overlapping, and attractive, if the cells are within an interaction distance, *r*_*max*_. First, we calculate the vector between cell centers: *x*_*ij*_ (*t*) = *x*_*j*_(*t*) − *x*_*i*_ (*t*). We then use Equation 3 to calculate the strength of the force between cells *i* and *j*, as a function of *x*_*ij*_ (*t*).

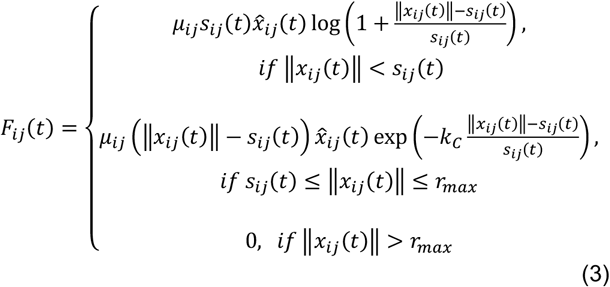

Here, *s*_*ij*_(*t*) is the sum of the radii of the two cells, *μ*_*ij*_ is the spring constant, 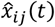 is the unit vector of *x*_*ij*_(*t*), *k*_*C*_ is the decay of the attractive force, and ‖*x*_*ij*_(*t*)‖ is the Euclidean norm of *x*_*ij*_(*t*). We note that we scale the forces based on a non-dimensionalized cell diameter of 1.

In addition, for each cell, we calculate its overlap with its neighboring cells: Δ*x*_*ij*_(*t*) = *s*_*ij*_(*t*) − ‖*x*_*ij*_ (*t*) ‖. We sum all the positive overlaps to yield Δ*x*_*i*_(*t*), which is the total overlap on a cell. If the total overlap exceeds a predefined threshold, we assume that the cell is compressed and is unable to proliferate, entering a quiescent state until it is uncompressed.

#### Forces exerted by the normal tissue

We assume that the tumor is surrounded by normal tissue cells. Besides impacting the dynamics of nutrients (described in the following section), we assume that this tissue exerts forces upon the tumor. Instead of modeling normal tissue cells individually, as this would greatly increase the computational time, we instead increase the damping coefficient, *η*, for cancer cells on the outer edge of the tumor. This slows their repulsion outward, representing their need to displace normal tissue.

### Diffusion

We model several diffusible factors, which exist in three categories: metabolic species, angiogenic factors, and chemotherapeutic agents. For metabolism, blood vessels supply oxygen and glucose, which cells uptake to produce ATP, and remove lactic acid (modeled as H^+^), a byproduct of glucose metabolism. For angiogenic factors, we model VEGF secretion by tumor cells. Lastly, we model a chemotherapeutic agent in a manner similar to nutrients, in that it diffuses out of the blood vessels to be taken up by cells.

The diffusion of these factors is modeled using partial-differential equations (PDEs). While cells are modeled using a continuous location, it is necessary to discretize the environment to numerically solve the PDEs. We solve them using a finite difference method. We note the assumption that diffusion is happening at a much faster rate than cellular processes, so that once the maximum fractional difference between diffusion step *n* and step *n* + 1 is less than 10^™4^, we assume the system has reached equilibrium and stop solving the PDEs until the next simulation step [28].

#### Metabolic species

Vessels (described in detail below) are modeled as point sources with a constant concentration of oxygen, glucose, and H+. We use the following equations (from Robertson-Tessi et al. [28]) to model cellular uptake of oxygen (O) and glucose (G) and the secretion of acid (H) (*ω*_*O*_, *ω*_*G*_, and *ω*_*H*_, respectively).

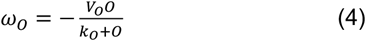

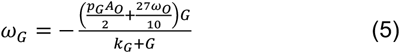

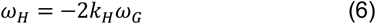

*V*_*O*_ is the uptake rate of oxygen under normoxic conditions and *A*_*O*_ is the ATP requirement of the cell, calculated as 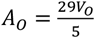. *k*_*O*_ and *k*_*G*_ are the half-maximal concentrations for oxygen and glucose uptake. As shown in Equation 5, glucose uptake is influenced by the oxygen uptake rate. Due to the Pasteur effect, as oxygen decreases, a cell will take up more glucose in order to meet its ATP demand. *p*_*G*_ is a scaling parameter that represents the Warburg effect, and is increased to represent increased aerobic glycolysis and the glycolytic shift by cancer cells. *K*_*H*_ is a buffering constant for acid production.

Besides the tumor cells that we discretely model, we assume that there is environmental uptake of nutrients by normal tissue. As the tumor grows, we assume that it replaces the normal tissue, so that locations where tumor cells have been no longer have environmental uptake. From this, we yield the full equation describing nutrient dynamics, with *i* representing the nutrient, *ω*_*i*_*d*_*x*_ representing the uptake/secretion by a discretely modeled cancer cell at location *x*, and *ω*_*i*_*n*_*x*_ representing uptake/secretion by normal tissue:

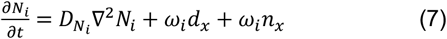

#### Angiogenic factor

Hypoxic cells secrete VEGF in order to recruit blood vessels, resupply nutrients, and clear waste products (i.e., lactic acid). VEGF degrades in the environment at a constant rate and is removed by vessels at a constant rate. Following Macklin et al. [30], we scale VEGF concentration to between 0 and 1. From this, we get Equation 8, with *λ*_*p*_ representing the production rate of VEGF, *d*_*h,x*_ representing the location of cells that produce VEGF, *λ*_*d,V*_ representing the degradation rate, and *λ*_*ves*_*ν*_*x*_ representing removal by the vasculature in locations where there are vessels.

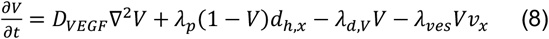

#### Chemotherapeutic agent

We model chemotherapy in a similar manner as nutrients, in that it diffuses out of blood vessels and is taken up by cells. We also assume degradation of the chemotherapeutic agent. The diffusion equation for chemotherapeutic agent is given in Equation 9, with *λ*_*C*_ as the uptake rate of the chemotherapeutic agent and *λ*_*d,C*_ as its degradation rate.

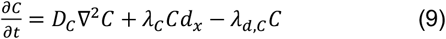

### Vasculature

We model vessels in a point fashion, initializing the environment following Robertson-Tessi et al. [28]. As the tumor grows, it exerts pressure on vessels. If the amount of overlap between a vessel and a tumor cell is above a threshold, we assume that the vessel collapses and is removed from simulation. As the tumor continues to grow, cells on the interior become hypoxic and secrete VEGF to recruit new blood vessels. To model this VEGF-mediated vascularization, at each timestep, the probability of recruiting a vessel at a location is equal to the VEGF concentration at that location multiplied by a base probability. This interplay between vessel collapse and vessel recruitment leads to intermittent regions of hypoxia and low pH that influence tumor development, as described in later sections.

A key feature of tumor vasculature is that it is unstable and not as capable of delivering blood to the tumor, making it more difficult for nutrients and chemotherapeutic agents to reach the tumor [3]. We model this feature with the introduction of a parameter for vascular health, *h*_*V*_. This value is set at 1.0 for vessels present at the start of simulation, however recruited vessels enter the environment with a lower value. We do not allow this value to change with time, except with the addition of anti-angiogenic therapy (explained in a later section). For interactions with tumor cells, we use this parameter to scale the spring constant, *μ*_*ij*_, between the vessel and tumor cells, making it easier for the tumor to collapse the recruited vessels. In order to reduce blood flow through these vessels, we use *h*_*V*_ to scale the rate of refresh nutrient concentrations at the blood vessels. At each diffusion time-step, *n*, when solving the PDEs, we update the concentration of each nutrient, *N*. Equation 10 displays how this method is discretely implemented in the model following the finite difference step to increase the concentration of nutrients at vessel locations. For healthy vessels, this maintains the concentration of nutrients in the vessels at blood concentration. For unstable vessels, this leads to a lower equilibrium concentration, reflecting the inability of recruited vasculature to properly deliver blood flow.

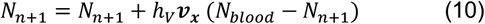

### Cell functions

#### Proliferation

We give each cell a probability of committing to the cell cycle at each time step. This probability starts at 1 and decreases in response to hypoxia (described below). If a cell has yet to commit to the cell cycle, it is in a migratory state. Once committing, we assume the cell stops migrating and grows until it can proliferate. We assume that cells proliferate away from current forces on the cell. Therefore, after calculating the current forces on a cell using the method described above, we place the daughter cell so that its center is one cell radius away from the mother cell’s center. These two cells then repel each other as described above.

#### Migration

If migratory, cells will migrate following the oxygen gradient with a biased random walk. In order to follow the gradient, we determine the cell’s *x-y* location based on its continuous position within the grid used to solve the PDEs. From this, we look at the oxygen concentrations in the Moore neighborhood of the cell and use these concentrations as probabilities for determining the direction of cell migration. After randomly selecting a direction, we calculate the angle between the cell’s current position and the chosen location and move the cell in that direction for a distance equal to the timestep duration multiplied by the migration speed. We bias the ability of cells to choose the direction with the highest oxygen concentration by multiplying the probability of the highest concentration by a scaling factor. This means that cells will closely follow the gradient, with some deviation that can be attributed to variations in the extracellular environment.

#### Metabolism

In order to carry out cell functions, cells need to produce enough ATP. We calculate ATP production using the above rates of oxygen and glucose uptake and Equation 11, taken from Robertson-Tessi et al. [28].

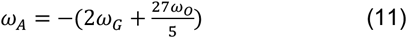

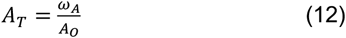

Here, *A*_*T*_ is the fraction of the ATP requirement met by the cell. We set ATP threshold values *A*_*Q*_ and *A*_*D*_. If *A*_*T*_ ≥ *A*_*Q*_, then the cell is in a proliferative state. If *A*_*Q*_ > *A*_*T*_ > *A*_*D*_, then the cell is not producing enough ATP to carry out proliferative functions and is in a quiescent state. If *A*_*T*_ ≤ *A*_*D*_, then the cell is not producing enough ATP to survive and becomes necrotic. Necrotic cells shrink and then are removed from the simulation.

#### Cell death

Cells also become necrotic and die if the local pH is below a certain threshold.

### Environmental effects

#### Hypoxia

When exposed to low concentrations of oxygen, tumor cells become hypoxic. This induces three changes. The first is that the cells secrete VEGF in order to recruit new blood vessels. The second is that the cells upregulate glucose transporters, modeled as an increase in *p*_*G*_, which increases glucose uptake and acid secretion. The third is that the probability of committing to the cell cycle decreases, making the cell more migratory. The increase in *p*_*G*_ and migratory ability is passed down to daughter cells. These hypoxia-induced changes are based on experimental evidence [10–13,16,17].

#### Low pH

Once the local pH falls below a certain threshold, a cell will become necrotic and die. However, we include the ability of cells to adapt to their changing environment by decreasing the threshold at which a cell will die [14,18]. We allow this effect to occur when the local pH is within 0.1 of the cell’s death threshold. We assume that there is an energy cost associated with this change and increase the cell cycle duration as cells become more resistant to low pH. At the highest level of acid resistance, the cell cycle duration is doubled [34]. This property is passed down to daughter cells. We give each cell a probability of committing to the cell cycle at each time step. This probability starts at 1 and decreases in response to hypoxia (described below). If a cell has yet to commit to the cell cycle, it is in a migratory state. Once committing, we assume the cell stops migrating and grows until it can proliferate. We assume that cells proliferate away from current forces on the cell. Therefore, after calculating the current forces on a cell using the method described above, we place the daughter cell so that its center is one cell radius away from the mother cell’s center. These two cells then repel each other as described above.

### Treatment

#### Anti-angiogenic therapy

We model anti-angiogenic treatment simply as turning off vessel recruitment to the tumor. We chose to use the complete cessation of angiogenesis because we wanted to examine tumor behavior at ideal therapeutic efficacy. We also allow anti-angiogenic treatment to normalize the vasculature [2,35], restoring vessel health, *h*_*V*_, back to that of non-recruited vessels, which makes it harder for the vessels to collapse and increases the rate at which they provide nutrients and chemotherapy. Thus, vascular normalization provides a temporary benefit to the tumor, but also increases the amount of chemotherapy delivered.

#### Chemotherapy

We model chemotherapy in a similar fashion as Pérez-Velázquez and Rejniak [36]. The concentration of the chemotherapeutic agent that cells are exposed to follows Equation 13:

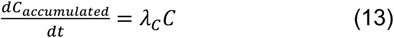

Subsequent damage caused by chemotherapy is modeled according to Equation 14:

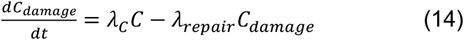

If the cell has been exposed to a certain amount of drug for a sufficient time period, then the cell gains tolerance to the drug, according to Equation 15. If *C*_*damage*_ > *C*_*tolerance*_, then the cell dies.

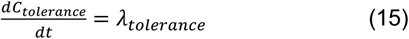

### Parameters

Most parameters are taken from the literature. Parameters not available in the literature were tuned to replicate known biological behavior. All model parameters are given in Supplementary Table 1.

## 3 RESULTS

### 3.1 Higher Rate of Angiogenesis Preserves Protective Outer Tumor Layer

We varied the rate of angiogenesis and the health of recruited vessels to see how these parameters impacted tumor growth and development. Simulations were run until the tumor reached a diameter that corresponded to a volume of 2 mm^3^.

We note that tumor diameter is calculated here as the largest distance between any two cancer cells, meaning that the simulation will end if cancer cells have sufficiently invaded into the surrounding tissue. In Figure 2, we show representative images of the final tumor state, with black cells representing the base phenotype and the colored cells representing the shift towards increased glycolysis (purple), increased acid resistance (blue), and increased migratory potential (yellow), with a more vibrant color representing a further shift. At the lower rate of vessel recruitment, we see that an increase in vascular health, while preventing some dissemination into the surrounding tissue, still has large numbers of cancer cells that are migrating away from the tumor. Additionally, most of the cells in these tumors adapted to harsher conditions, making the tumor more invasive. At a higher rate of vessel recruitment, tumors retain an outer layer of cells in the initial phenotype. Here, the outer layer of tumor cells was not exposed to harsher environmental conditions, thus there was no pressure to adapt. With this layer acting as a barrier and the sufficient supply of oxygen into the center of the tumor by recruited vessels (Fig S1), the more invasive cells are retained within the tumor core and fail to spread outward.

**Figure 2.**
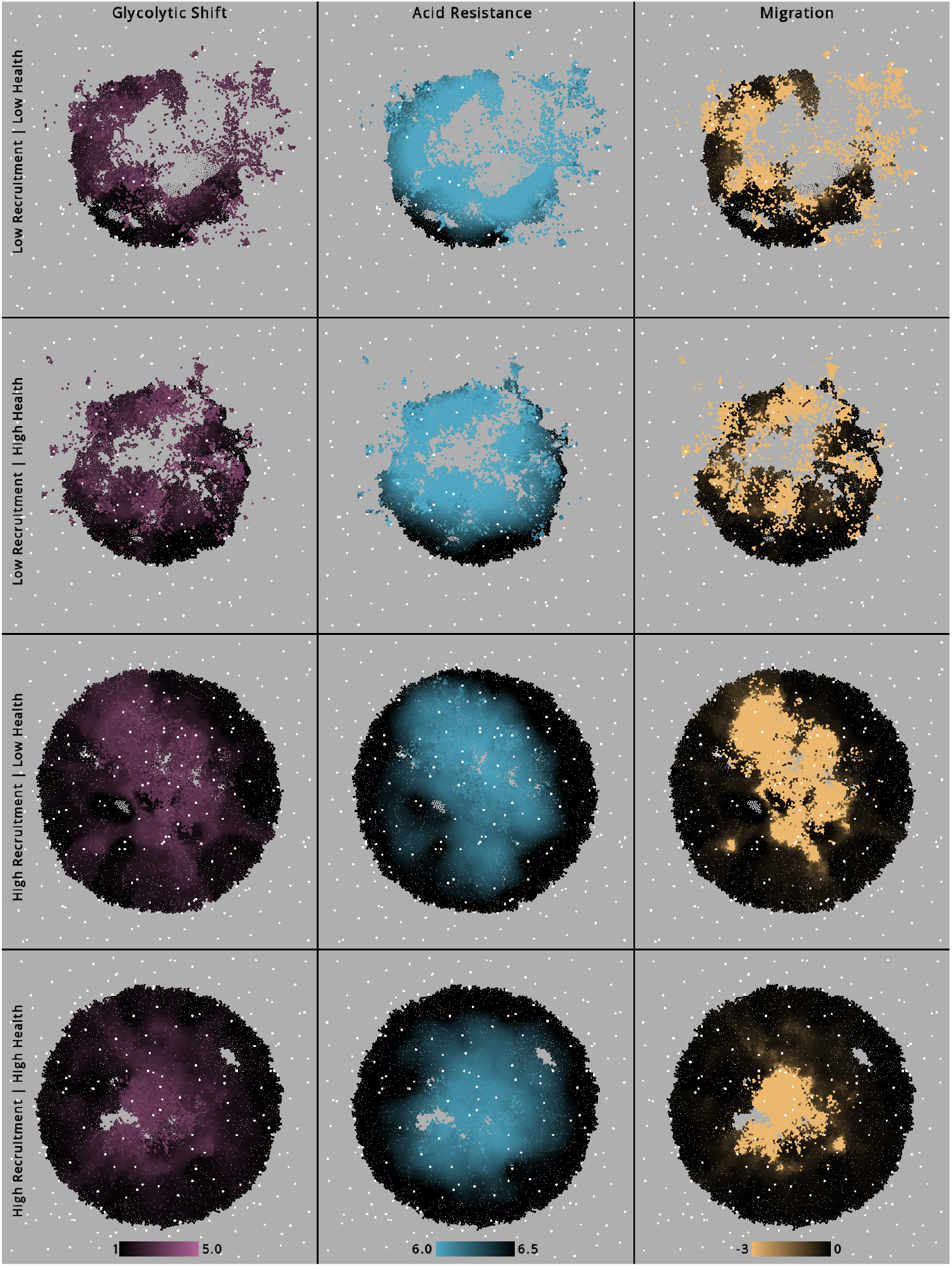
Predicted tumor cell properties without treatment. Spatial layout of representative final tumor states at different rates of vessel recruitment (1.58 × 10^−6^ and 3.16 × 10^−6^(*hr* × *μm*^2^) ^−1^) and different levels of recruited vessel health (0.25 and 0.5). Phenotypic properties are shown in the three columns, with black cells representing the base phenotype and colored cells representing increases in glycolysis (purple), acid resistance (blue), and migratory potential (yellow) as the log_10_ of the probability of committing to the cell cycle. Vessels are shown in white. Color bars at the bottom of each column show the range of phenotypic properties for this figure and the other spatial figures that we present.

### 3.2 Anti-Angiogenic Treatment Promotes Tumor Evolution and Local Invasion

We then looked at how anti-angiogenic treatment impacts tumor shape and the distribution of tumor cells with the three properties of interest (glycolytic shift, acid resistance, and migratory potential). Here, and for the rest of the study, we focused our simulations on one combination of vessel recruitment rate and vascular health, higher recruitment (3.16 × 10^−6^(*hr* × *μm*^2^)^−1^) and lower health (0.25). Figure 3 shows a comparison of representative tumor simulations without treatment (left) and with treatment (right). Here, we simulated the no-treatment case until the tumor reached a diameter corresponding to a volume of 2 mm3, as described above. We then simulated the treatment case for the same time duration as the no-treatment case. Treatment was started when the tumor reached 1 mm^3^. While anti-angiogenic treatment succeeded in greatly reducing the number of cancer cells, we see much further dissemination into the surrounding tissue along with an absence of cells in the tumor core where necrosis occurred. This is consistent with clinical and in vivo observations that anti-angiogenic treatment promotes local invasion as cancer cells are forced to search for a more habitable environment [7–9]. We also see that most of the vasculature in the simulation environment was collapsed by the invading tumor.

**Figure 3.**
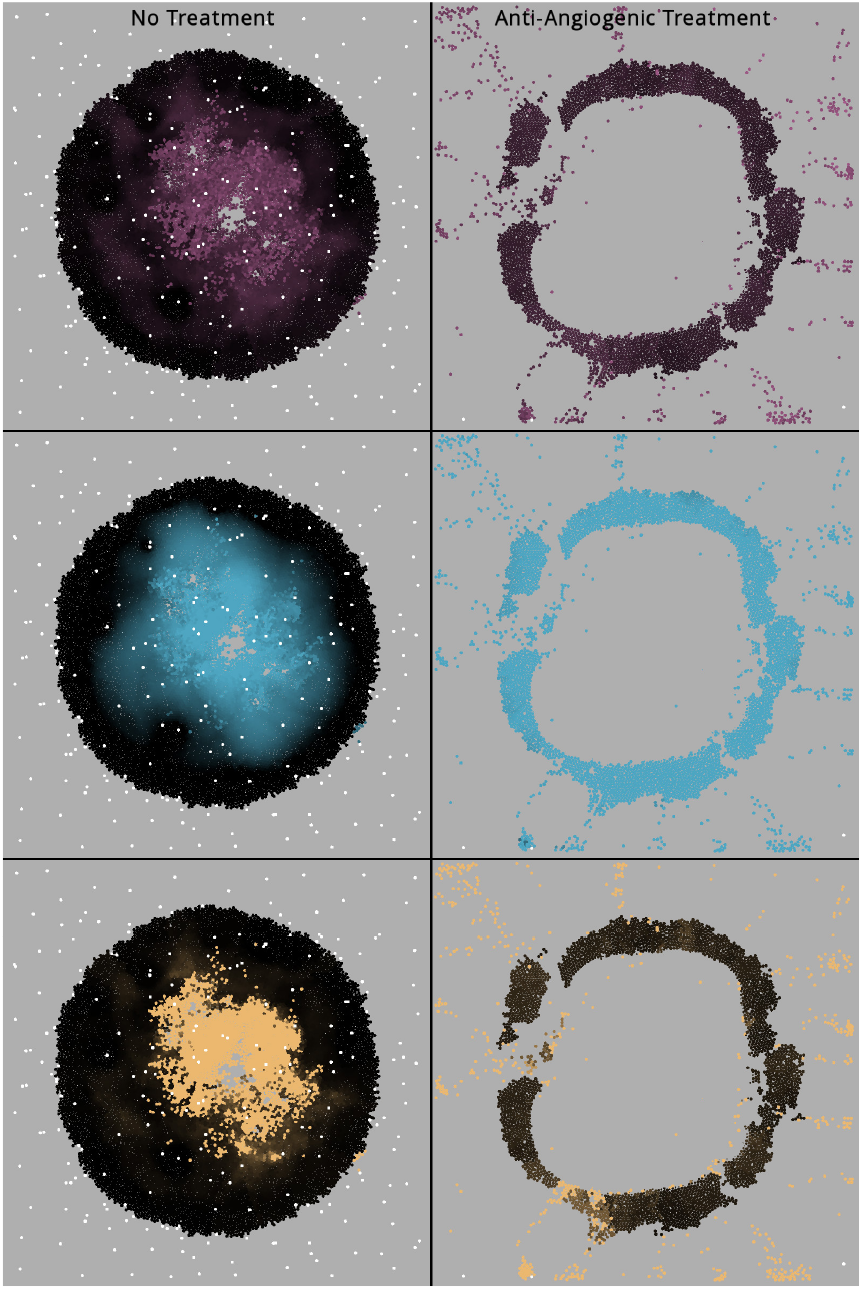
Tumor cells’ response to anti-angiogenic treatment. Spatial layout of sample final tumor states without treatment (left) and with anti-angiogenic treatment (right). Cancer cell phenotype is shown in the three rows, with black cells representing the base phenotype and colored cells representing increases in glycolysis (purple), acid resistance (blue), and migratory potential (yellow). Vessels are shown in white.

In Figure 4 we show frequency distributions of cellular properties for 50 simulation replicates, comparing simulations with anti-angiogenic treatment (red) and the no-treatment case (black). This provides quantitative evidence of the qualitative observations from the spatial layout: there is a clear shift in cell location away from the center of the simulation environment due to anti-angiogenic treatment (Figure 4A). In addition, cells display increased glycolysis (Figure 4B), a lower pH threshold (Figure 4C), and a lower probability of committing to the cell cycle, thus being more migratory (Figure 4D). Altogether, the simulations indicate the way in which anti-angiogenic therapy promotes the phenotypic shift of tumor cells, making them more tolerant to harsher microenvironmental conditions.

**Figure 4.**
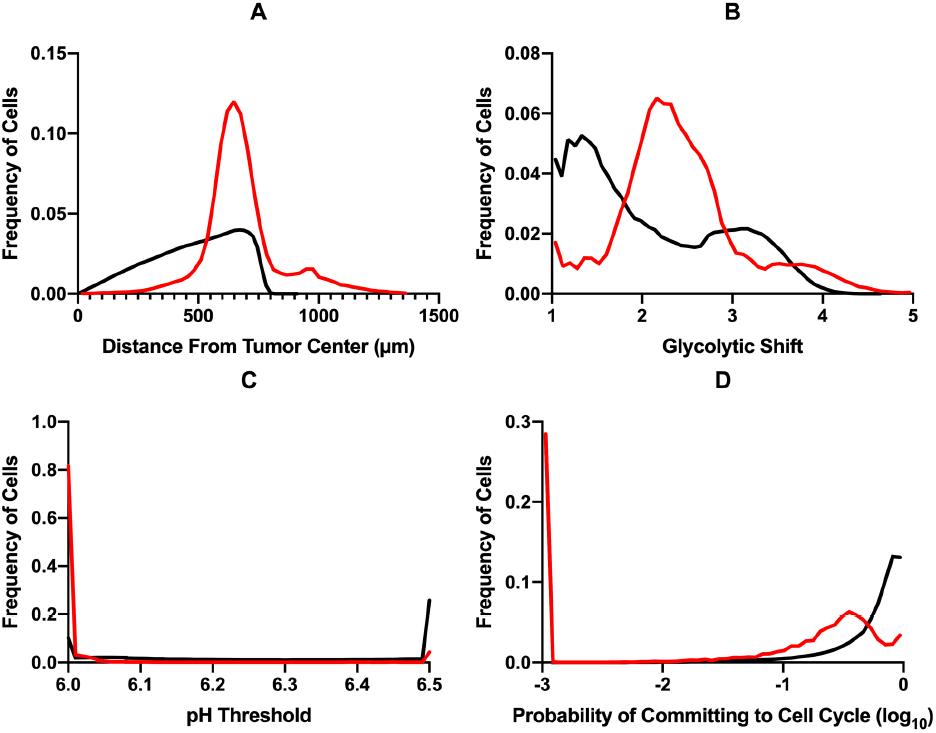
Distributions of cell properties. Frequency distributions for 50 simulation replicates showing cell distance from the tumor center in micrometers (A), increase in the value of *p*_*G*_ (B), decrease in the pH value that induces cell death (C), and decrease in the probability of committing to the cell cycle, meaning an increase in cell migration (D). Distributions without treatment are shown in black while distributions with anti-angiogenic treatment are in red. Note that at low values of the x-axis for panel (D), the red and black curves overlap.

### 3.3 Anti-Angiogenic Therapy Limits Tumor Growth After Chemotherapy

We next examined whether augmenting chemotherapy with anti-angiogenic therapy can improve the chemotherapeutic effects. We simulated a chemotherapeutic agent that diffuses out of the blood vessels. Cancer cells died once accumulating sufficient damage, based on the concentration of the chemotherapeutic agent. In addition, cells acquired tolerance to chemotherapy at a constant rate upon sufficient exposure to the drug. Both therapies were started once the tumor diameter corresponded to 1 mm^3^ and ended either when the diameter corresponded to 2 mm^3^ or simulation time reached 100 days, whichever occurred first. Anti-angiogenic therapy was simulated as constantly on, whereas chemotherapy was cycled on for five days and off for ten until the simulation ended. This combination of continuous anti-angiogenic therapy and intermittent chemotherapy has been used in pre-clinical studies [22].

Time courses for cancer cell numbers with chemotherapy alone (blue) and chemotherapy augmented by anti-angiogenic therapy (red) are shown in Figure 5. To investigate how the response to combination therapy changes as a function of drug resistance, we varied the rate at which cancer cells became resistant to the chemotherapeutic agent. At a lower rate of acquired resistance (Figure 5 Ai), we see that the number of tumor cells is greatly reduced and maintained at a low number, with simulations ending at 100 days instead of meeting the stopping criteria based on tumor diameter, indicating that the cancer cells have not invaded into the surrounding tissue. At a higher rate of acquiring resistance (Figure 5 Bi), we see that with chemotherapy alone, the tumor begins to regrow at a much faster rate than with the addition of anti-angiogenic therapy. At the highest rate of resistance, we see a rapid regrowth of both populations (Figure 5 Ci), however the final cell population is much smaller with combination therapy. These simulations end before 100 days due to migration away from the tumor. This is shown in Figure 5ii, whereas resistance increases, the distribution of cell distance from the center increases for combination therapy. For chemotherapy alone, as resistance increases, the distribution starts to resemble that of no treatment at all (Figure 4A). Interestingly, even at the highest rate of resistance, the distance distribution for combination therapy is not as extreme as that for anti-angiogenic therapy alone (dotted black line, Figure 5iii). This is likely due to the continued killing effect of chemotherapy as migratory cells approach blood vessels.

**Figure 5.**
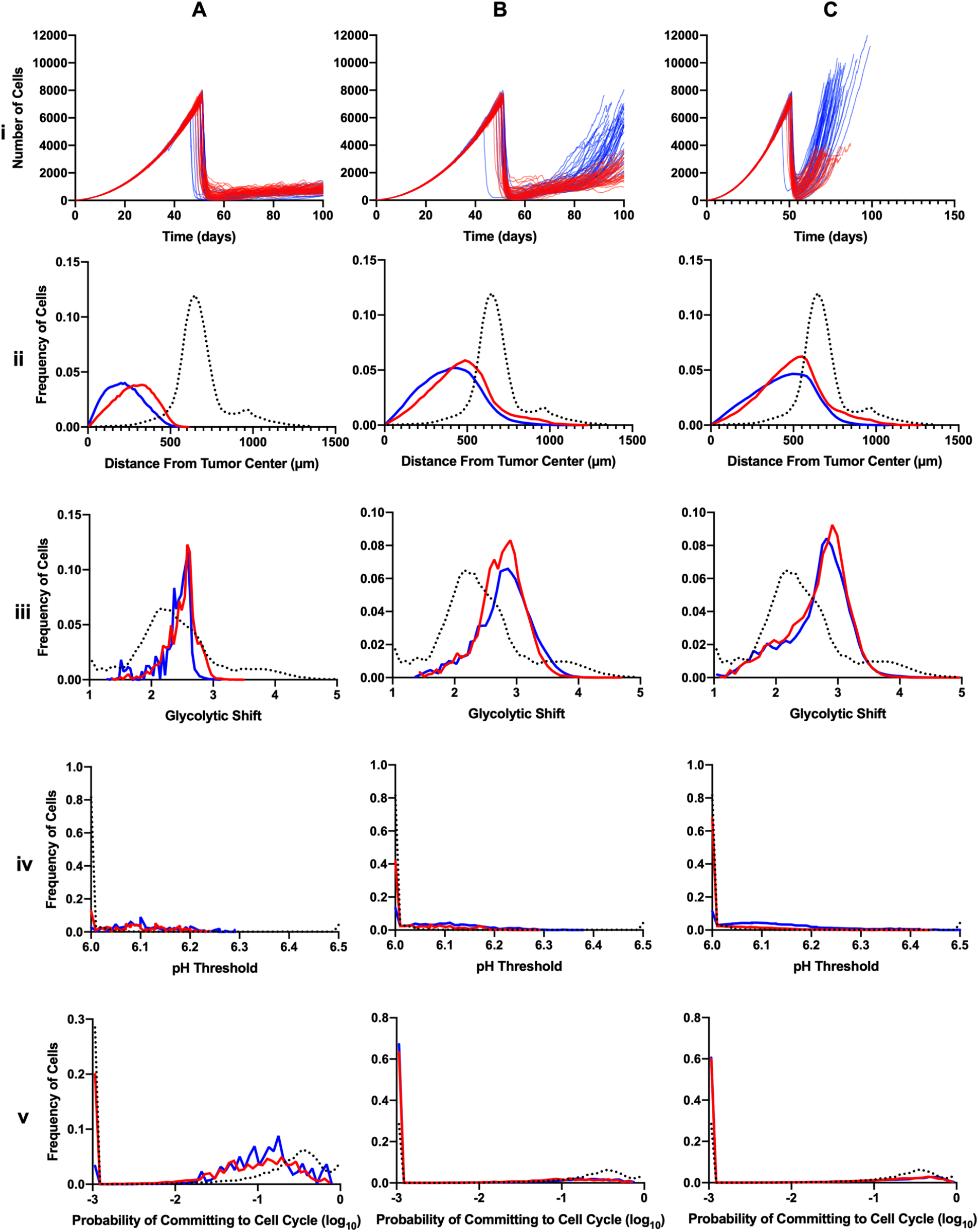
Effect of acquired tolerance and single-agent or combined therapy on tumor properties. Time-courses (i) for different rates of acquiring tolerance to chemotherapy: 10^™5^ (A), 10^™4.5^ (B), and 10^™4^ (C). Frequency distributions for the distance from tumor center (ii), increase in *p*_*G*_ (iii), pH value that induces cell death (iv), and probability of committing to the cell cycle (v). Simulations are shown for 50 replicates where the tumor was not removed by therapy for chemotherapy alone (blue), chemotherapy augmented with anti-angiogenic therapy (red), and anti-angiogenic therapy (black dotted line, from Figure 4).

Interestingly, we see that the addition of anti-angiogenic therapy has little influence on the phenotypic distributions when compared to chemotherapy alone (Figure 5iii-v). What is also of note when looking at glycolytic shift is that, while these distributions are shifted further to the right than with anti-angiogenic therapy alone, we see that a subpopulation of the cells from anti-angiogenic therapy alone reach a higher shift than both chemotherapy and combination therapy. We also see that a greater portion of the cells with anti-angiogenic therapy alone are at the lowest pH for acid-induce cell death, however as resistance increases, the cells exposed to combination therapy achieve a similar distribution.

In Figure 6, we show representative final tumor states for each rate of resistance from Figure 5, comparing the effects of chemotherapy without and with anti-angiogenic treatment. We note that in this figure we only show cell location, and phenotypic properties are shown in Fig S2 and S3. From these figures we see that, most prominently at the lowest rate of acquiring resistance, chemotherapy alone yields tumor regrowth that has distinct clusters of cells, instead of the fully dense tumors predicted from simulations without any treatment (Figure 2). With the addition of anti-angiogenic therapy, tumor growth resembles the cases with only anti-angiogenic therapy, with few cells towards the tumor center and groups of cells migrating into the surrounding tissue.

**Figure 6.**
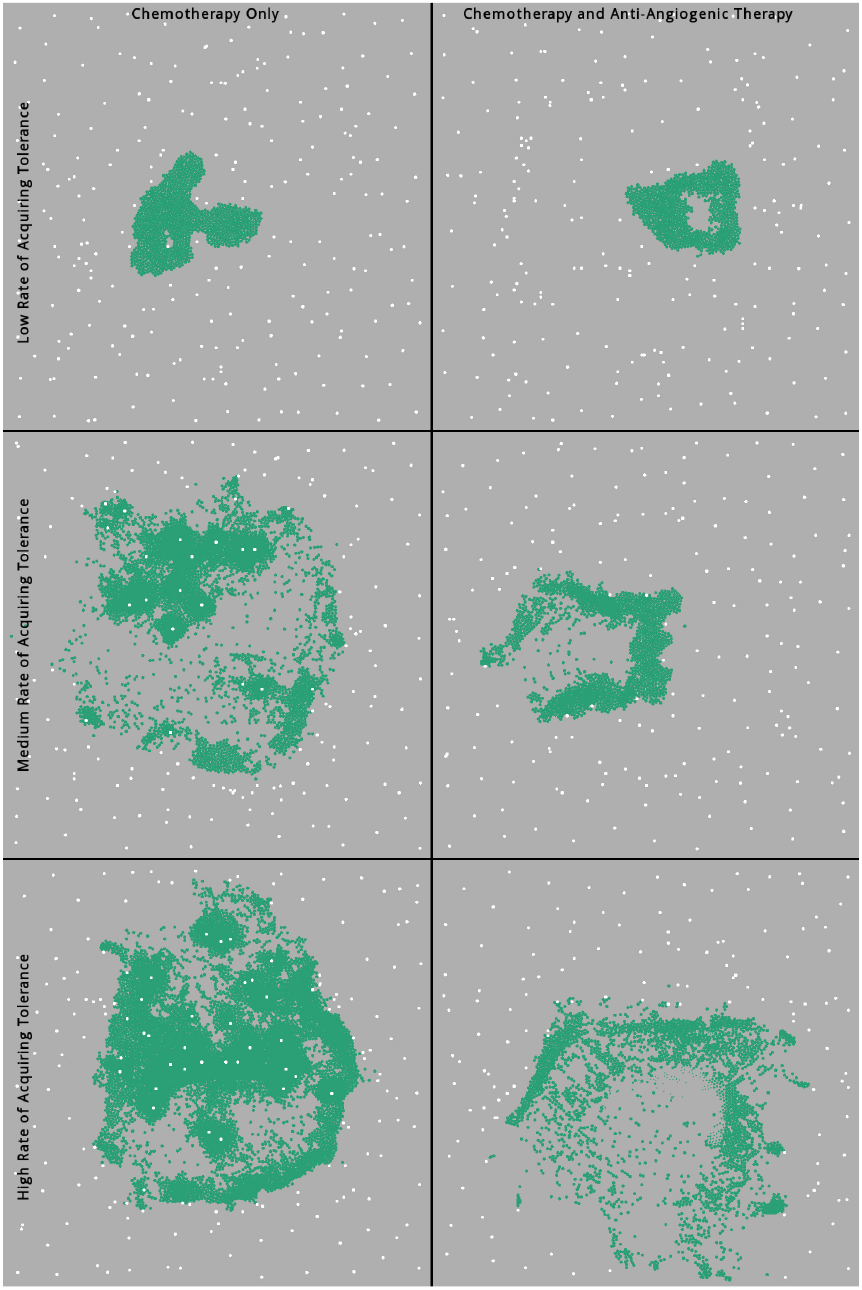
Effect of acquired tolerance and single-agent or combined therapy on final tumor states. Spatial layout for representative final tumor states with chemotherapy alone (left) and chemotherapy with anti-angiogenic therapy (right), for different rates of acquiring resistance. Vessels are shown in white and cancer cells are shown in green.

## 4 DISCUSSION

In this study, we highlight how the use of anti-angiogenic therapies can lead to a more invasive tumor that is better adapted to harsher microenvironmental conditions. Our results agree with previous *in vivo* and clinical studies that find that local invasion is increased by anti-angiogenic therapy [7–9]. By using a computational model, we are able to further explore this phenomenon, examining how the tumor not only evolves spatially, but adapts phenotypically to the environmental pressure exerted on it by treatment. The metabolic changes predicted by the model, increased glycolysis and resistance to acidic pH, are associated with more aggressive tumors, allowing these cells to better invade into surrounding tissue.

Here, we found that anti-angiogenic therapy alone prompts cancer cells that have developed in the harsh microenvironment of the tumor core to migrate out into the surrounding tissue in search of more habitable regions. Compared to simulations without any treatment, we find that preventing angiogenesis causes further phenotypic shifts towards properties that are induced by hypoxia and low pH. From this, we see that while anti-angiogenesis alone can reduce the number of cancer cells, it also has the potential to advance tumor development by creating a microenvironment that selects for more robust and invasive cancer cells. This highlights how care must be taken to avoid Darwinian selection of cancer cells that are more fit for harsher environments [37].

Despite these negative effects of introducing anti-angiogenic treatment, the model predicts that anti-angiogenic treatment provides some benefits when combined with chemotherapy. The combination of these two strategies has been shown to be effective in some cancers [2,3,38]. Indeed, we found that including anti-angiogenic therapy is able to slow tumor regrowth following chemotherapy; however, there was little difference in phenotypic shift between chemotherapy alone and the addition of anti-angiogenic therapy.

Our model is limited in some ways, which can be addressed in future studies. Here, we kept the rates of phenotypic shift constant, choosing instead to focus on the effects of therapy. We can expand the model by examining how these rates impact tumor development with and without the addition of anti-angiogenic therapy. Additionally, our treatment of normal tissue is limited, represented only by an increase in the damping coefficient. We can extend our modeling of normal tissue to include more explicit tissue forces. We can also expand this section of the model to include destruction of normal cells and extracellular matrix by lowered pH, thus increasing the model’s detail on how acidity enables tumor invasion.

## 5 CONCLUSIONS

Here, we find that anti-angiogenic therapy promotes tumor invasion by exposing the tumor to a harsher microenvironment that forces the cancer cells to adapt in order to survive. These cells exhibited further shifts in aerobic glycolysis, acid resistance, and migratory potential compared to simulations without any treatment. Additionally, we find that augmenting chemotherapy with anti-angiogenic therapy can prolong tumor regrowth following the acquisition of resistance. Overall, our model generates quantitative and mechanistic insights into experimental and clinical observations following anti-angiogenic treatment.

## ACKNOWLEDGEMENTS

The authors thank members of the Finley research group for critical comments and suggestions. This work was supported by the American Cancer Society (130432-RSG-17-133-01-CSM to SDF) and the USC Provost’s PhD Fellowship (CGC).

## SUPPORTING INFORMATION

**File S1**. Supplementary figures and table (pdf file).

## REFERENCES

[1] D. Hanahan and R.A. Weinberg, Hallmarks of cancer: the next generation, Cell. 144 (2011) 646–674.

[2] R. Lugano, M. Ramachandran, and A. Dimberg, Tumor angiogenesis: causes, consequences, challenges and opportunities, Cellular and Molecular Life Sciences. 77 (2020) 1745–1770.

[3] A.M. Abdalla, L. Xiao, M.W. Ullah, M. Yu, C. Ouyang, and G. Yang, Current challenges of cancer anti-angiogenic therapy and the promise of nanotherapeutics, Theranostics. 8 (2018) 533.

[4] E. Maj, D. Papiernik, and J. Wietrzyk, Antiangiogenic cancer treatment: The great discovery and greater complexity, International Journal of Oncology. 49 (2016) 1773–1784.

[5] G. Lupo, N. Caporarello, M. Olivieri, M. Cristaldi, C. Motta, V. Bramanti, R. Avola, M. Salmeri, F. Nicoletti, and C.D. Anfuso, Anti-angiogenic therapy in cancer: downsides and new pivots for precision medicine, Frontiers in Pharmacology. 7 (2017) 519.

[6] N.S. Vasudev and A.R. Reynolds, Anti-angiogenic therapy for cancer: current progress, unresolved questions and future directions, Angiogenesis. 17 (2014) 471–494.

[7] J.M. Ebos, C.R. Lee, W. Cruz-Munoz, G.A. Bjarnason, J.G. Christensen, and R.S. Kerbel, Accelerated metastasis after short-term treatment with a potent inhibitor of tumor angiogenesis, Cancer Cell. 15 (2009) 232–239.

[8] M. Pàez-Ribes, E. Allen, J. Hudock, T. Takeda, H. Okuyama, F. Viñals, M. Inoue, G. Bergers, D. Hanahan, and O. Casanovas, Antiangiogenic therapy elicits malignant progression of tumors to increased local invasion and distant metastasis, Cancer Cell. 15 (2009) 220–231.

[9] D. Wang, C. Tan, F. Xiao, L. Zou, L. Wang, H. Yang, and W. Zhang, The “inherent vice” in the anti-angiogenic theory may cause the highly metastatic cancer to spread more aggressively, Scientific Reports. 7 (2017) 1–14.

[10] W. Al Tameemi, T.P. Dale, R.M.K. Al-Jumaily, and N.R. Forsyth, Hypoxia-modified cancer cell metabolism, Frontiers in Cell and Developmental Biology. 7 (2019) 4.

[11] I.F. Robey, A.D. Lien, S.J. Welsh, B.K. Baggett, and R.J. Gillies, Hypoxia-inducible factor-1α and the glycolytic phenotype in tumors, Neoplasia. 7 (2005) 324–330.

[12] K. Eales, K. Hollinshead, and D. Tennant, Hypoxia and metabolic adaptation of cancer cells, Oncogenesis. 5 (2016) e190–e190.

[13] A.M. Weljie and F.R. Jirik, Hypoxia-induced metabolic shifts in cancer cells: moving beyond the Warburg effect, The International Journal of Biochemistry & Cell Biology. 43 (2011) 981–989.

[14] R.A. Gatenby, E.T. Gawlinski, A.F. Gmitro, B. Kaylor, and R.J. Gillies, Acid-mediated tumor invasion: a multidisciplinary study, Cancer Research. 66 (2006) 5216–5223.

[15] V. Estrella, T. Chen, M. Lloyd, J. Wojtkowiak, H.H. Cornnell, A. Ibrahim-Hashim, K. Bailey, Y. Balagurunathan, J.M. Rothberg, B.F. Sloane, J. Johnson, R.A. Gatenby, and R.J. Gillies, Acidity generated by the tumor microenvironment drives local invasion, Cancer Research. 73 (2013) 1524–1535.

[16] N.H. Barrak, M.A. Khajah, and Y.A. Luqmani, Hypoxic environment may enhance migration/penetration of endocrine resistant MCF7-derived breast cancer cells through monolayers of other non-invasive cancer cells in vitro, Scientific Reports. 10 (2020) 1–14.

[17] C.S. Daly, A. Flemban, M. Shafei, M.E. Conway, D. Qualtrough, and S.J. Dean, Hypoxia modulates the stem cell population and induces EMT in the MCF-10A breast epithelial cell line, Oncology Reports. 39 (2018) 483–490.

[18] M. Damaghi and R. Gillies, Phenotypic changes of acid-adapted cancer cells push them toward aggressiveness in their evolution in the tumor microenvironment, Cell Cycle. 16 (2017) 1739–1743.

[19] Q. Wu and S.D. Finley, Predictive model identifies strategies to enhance TSP1-mediated apoptosis signaling, Cell Communication and Signaling. 15 (2017) 1–17.

[20] Q. Wu and S.D. Finley, Mathematical Model Predicts Effective Strategies to Inhibit VEGF-eNOS Signaling, Journal of Clinical Medicine. 9 (2020) 1255.

[21] D. Li and S.D. Finley, Exploring the extracellular regulation of the tumor angiogenic interaction network using a systems biology model, Frontiers in Physiology. 10 (2019) 823.

[22] D. Li and S.D. Finley, Mechanistic insights into the heterogeneous response to anti-VEGF treatment in tumors, Computational and Systems Oncology. 1 (2021) e1013.

[23] G. Vilanova, I. Colominas, and H. Gomez, A mathematical model of tumour angiogenesis: growth, regression and regrowth, Journal of The Royal Society Interface. 14 (2017) 20160918.

[24] J. Xu, G. Vilanova, and H. Gomez, A mathematical model coupling tumor growth and angiogenesis, PloS One. 11 (2016) e0149422.

[25] M.M. Olsen and H.T. Siegelmann, Multiscale agent-based model of tumor angiogenesis, Procedia Computer Science. 18 (2013) 1016–1025.

[26] K.-A. Norton, K. Jin, and A.S. Popel, Modeling triple-negative breast cancer heterogeneity: Effects of stromal macrophages, fibroblasts and tumor vasculature, Journal of Theoretical Biology. 452 (2018) 56–68.

[27] X. Sun, L. Zhang, H. Tan, J. Bao, C. Strouthos, and X. Zhou, Multi-scale agent-based brain cancer modeling and prediction of TKI treatment response: incorporating EGFR signaling pathway and angiogenesis, BMC Bioinformatics. 13 (2012) 1–14.

[28] M. Robertson-Tessi, R.J. Gillies, R.A. Gatenby, and A.R. Anderson, Impact of metabolic heterogeneity on tumor growth, invasion, and treatment outcomes, Cancer Research. 75 (2015) 1567–1579.

[29] M. Wu, H.B. Frieboes, S.R. McDougall, M.A. Chaplain, V. Cristini, and J. Lowengrub, The effect of interstitial pressure on tumor growth: coupling with the blood and lymphatic vascular systems, Journal of Theoretical Biology. 320 (2013) 131–151.

[30] P. Macklin, S. McDougall, A.R. Anderson, M.A. Chaplain, V. Cristini, and J. Lowengrub, Multiscale modelling and nonlinear simulation of vascular tumour growth, Journal of Mathematical Biology. 58 (2009) 765–798.

[31] M. Shamsi, M. Saghafian, M. Dejam, and A. Sanati-Nezhad, Mathematical modeling of the function of Warburg effect in tumor microenvironment, Scientific Reports. 8 (2018) 1–13.

[32] H. Perfahl, H.M. Byrne, T. Chen, V. Estrella, T. Alarcón, A. Lapin, R.A. Gatenby, R.J. Gillies, M.C. Lloyd, P.K. Maini, M. Reuss, and M.R. Owen, Multiscale modelling of vascular tumour growth in 3D: the roles of domain size and boundary conditions, PloS One. 6 (2011) e14790.

[33] J.M. Osborne, A.G. Fletcher, J.M. Pitt-Francis, P.K. Maini, and D.J. Gavaghan, Comparing individual-based approaches to modelling the self-organization of multicellular tissues, PLoS Computational Biology. 13 (2017) e1005387.

[34] A. Ibrahim-Hashim, M. Robertson-Tessi, P.M. Enriquez-Navas, M. Damaghi, Y. Balagurunathan, J.W. Wojtkowiak, S. Russell, K. Yoonseok, M.C. Lloyd, M.M. Bui, J.S. Brown, A.R.A. Anderson, R.J. Gillies, and R.A. Gatenby, Defining cancer subpopulations by adaptive strategies rather than molecular properties provides novel insights into intratumoral evolution, Cancer Research. 77 (2017) 2242–2254.

[35] R.K. Jain, Normalization of tumor vasculature: an emerging concept in antiangiogenic therapy, Science. 307 (2005) 58–62.

[36] J. Pérez-Velázquez and K.A. Rejniak, Drug-induced resistance in micrometastases: analysis of spatio-temporal cell lineages, Frontiers in Physiology. 11 (2020) 319.

[37] E. Kim, J.S. Brown, Z. Eroglu, and A.R. Anderson, Adaptive Therapy for Metastatic Melanoma: Predictions from Patient Calibrated Mathematical Models, Cancers. 13 (2021) 823.

[38] J. Ma and D.J. Waxman, Combination of antiangiogenesis with chemotherapy for more effective cancer treatment, Molecular Cancer Therapeutics. 7 (2008) 3670–3684.

