## Supplementary File for "Multiscale modeling of tumor adaption and invasion following anti-angiogenic therapy"

### Supplementary Information

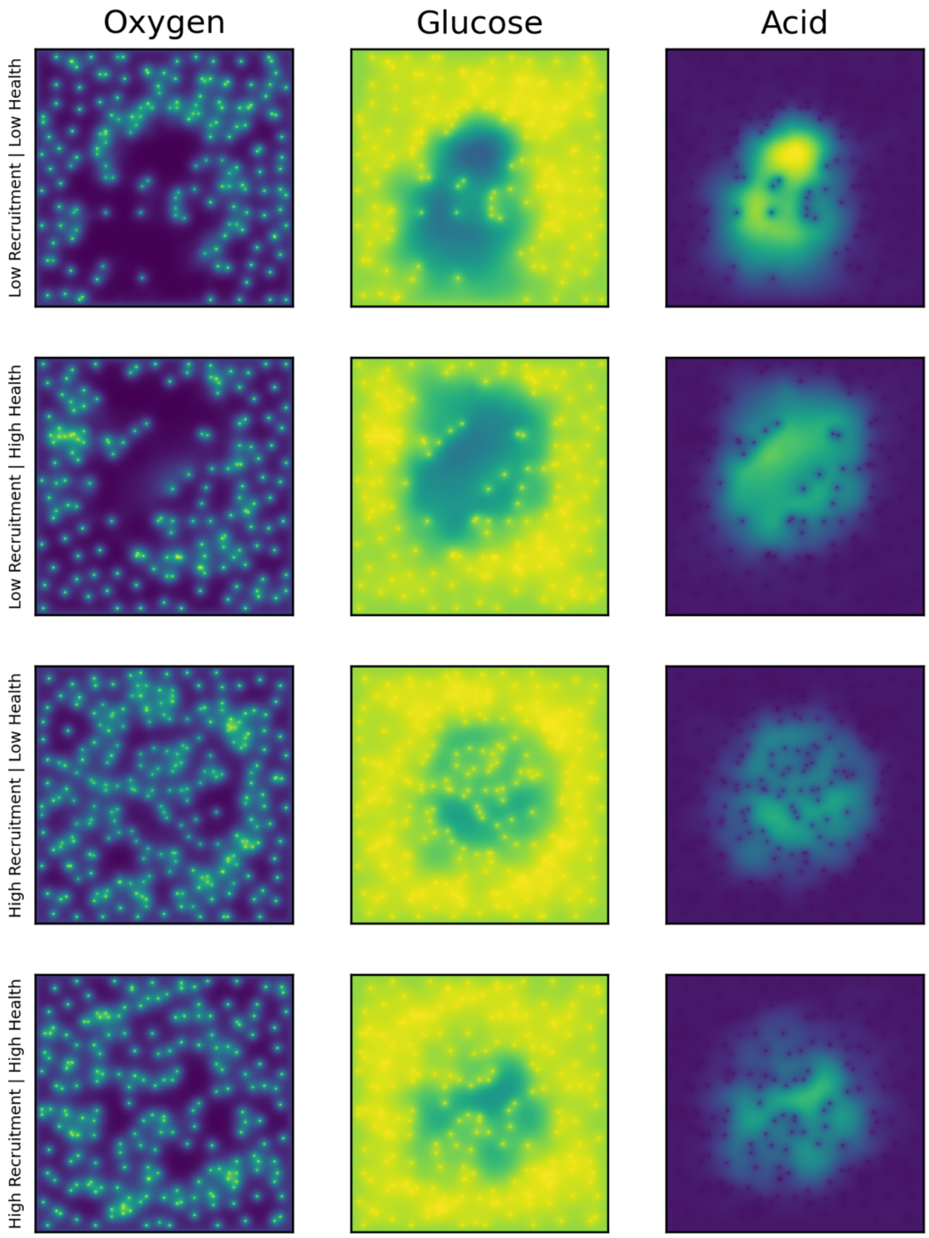


**Figure S1. Concentration gradients for sample simulations without treatment.** Concentration gradients for oxygen, glucose, and H+ for a sample simulation for the four combinations of vessel recruitment rate and vessel health. Yellow represents the highest value and purple the lowest value.

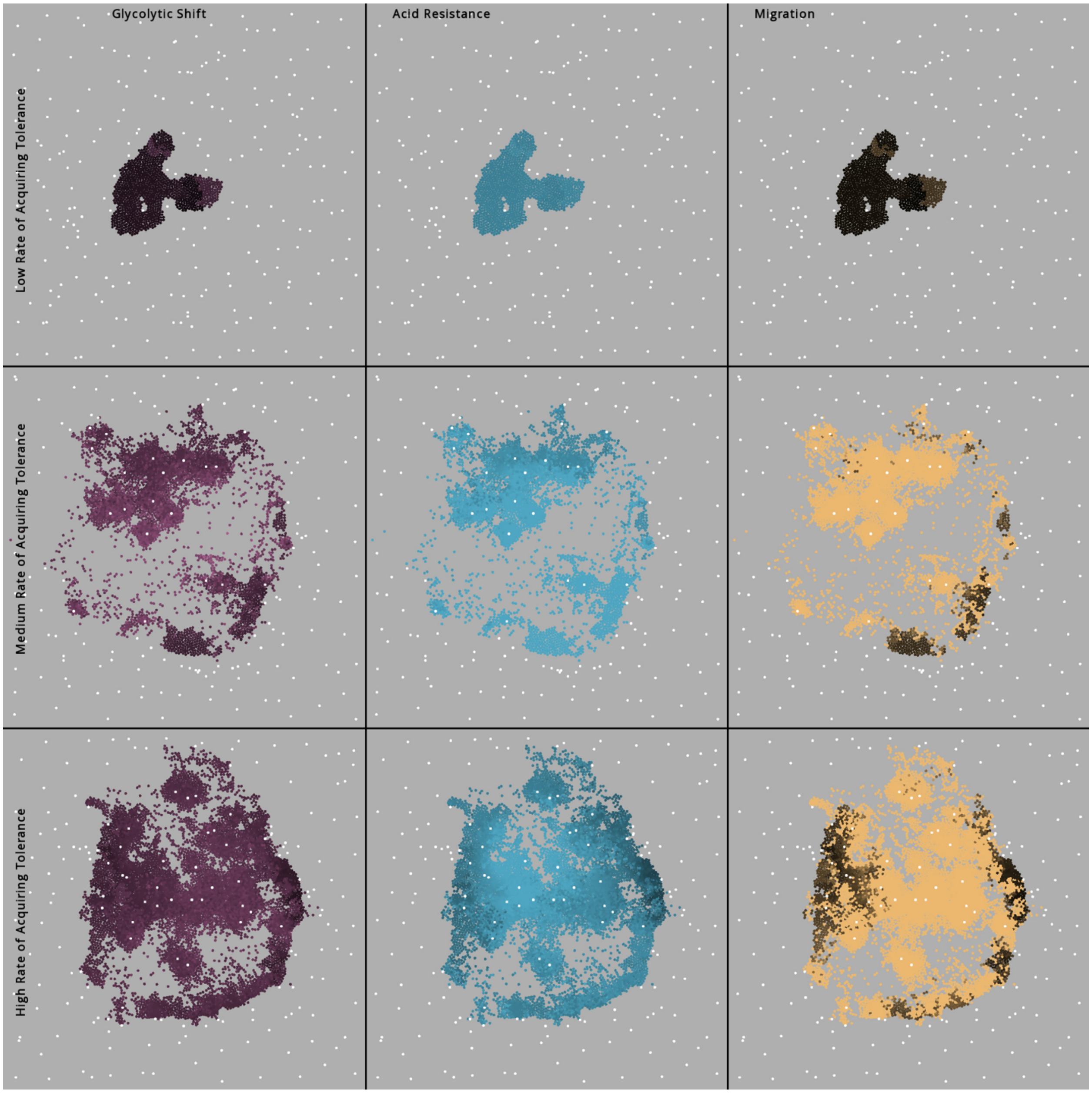


**Figure S2. Spatial layout of tumor phenotypes after chemotherapy.** Spatial layout of the phenotypic properties following chemotherapy from Figure 6.

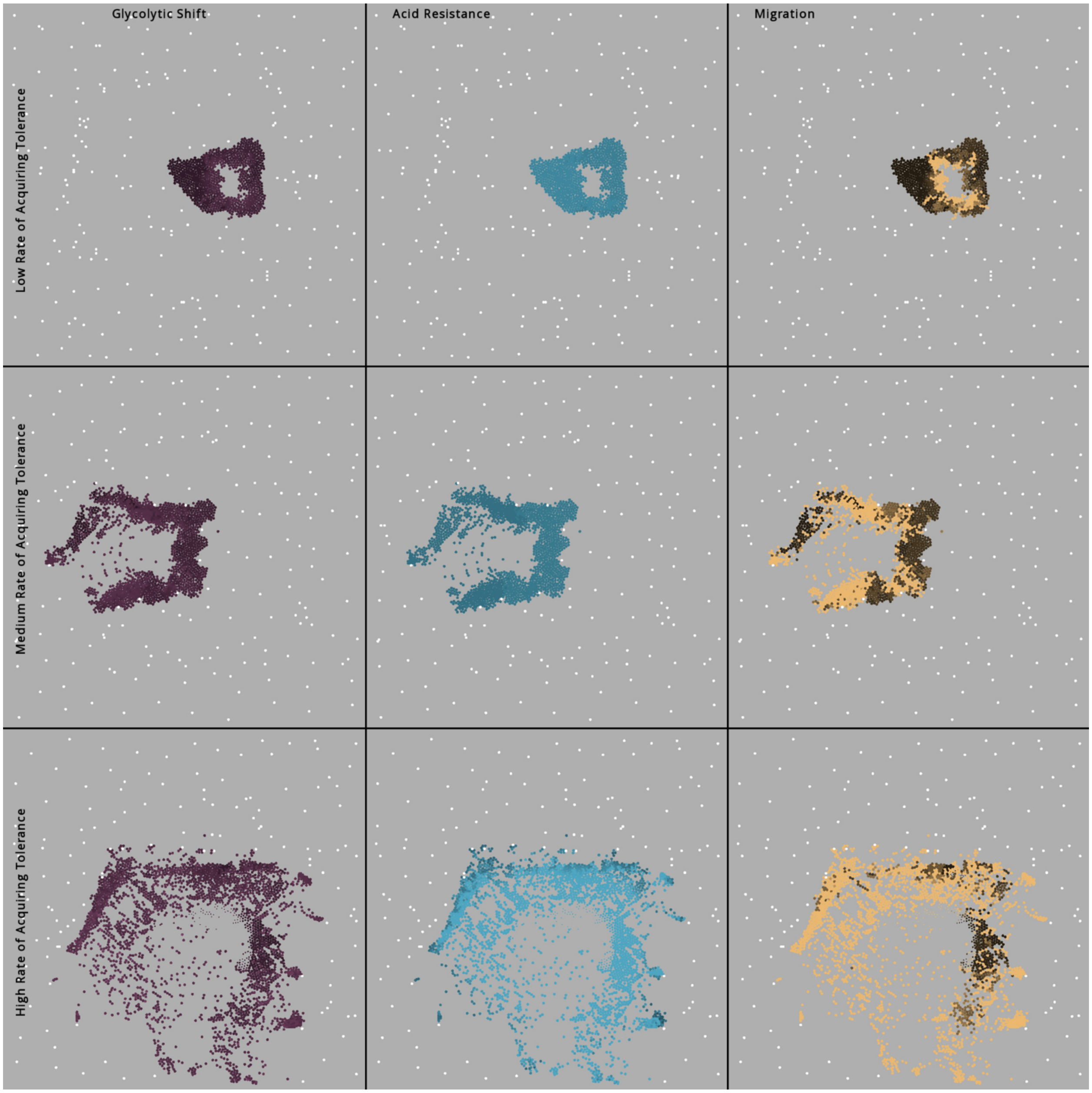


**Figure S3. Spatial layout of tumor phenotypes after chemotherapy and anti-angiogenic therapy.** Spatial layout of the phenotypic properties following combination therapy from Figure 6.

**Supplementary Table 1. Model parameters**

*Cancer Cell Parameters*

| Symbol | Description | Value | Reference |
|---|---|---|---|
| - | Cell diameter | 15 µm | [1] |
| $\boldsymbol{\mu}$ | Spring constant | 50 | [2] |
| $\boldsymbol{k_c}$ | Decay of attractive force | 10 | * |
| $\boldsymbol{\eta}$ | Damping coefficient | 10 | * |
| - | Increase factor for damping coefficient in normal tissue | 10 | * |
| - | Maximum overlap | 0.2 cell diameters | * |
| $\boldsymbol{V_O}$ | Maximum oxygen uptake rate | 0.012 mmol/L/s | [3] |
| $\boldsymbol{k_O}$ | Half-max oxygen concentration | 0.005 mmol/L | [3] |
| $\boldsymbol{k_G}$ | Half-max glucose concentration | 0.04 mmol/L | [3] |
| $\boldsymbol{p_g}$ | Initial glycolytic shift | 1.0 | [3] |
| $\boldsymbol{k_H}$ | Proton buffering coefficient | 2.5e-4 | [3] |
| - | Initial pH threshold for cell death | 6.5 | [3] |
| - | Rate of $p_G$ shift when hypoxic | 0.005 /hr | * |
| - | Maximum $p_G$ | 5.0 | * |
| - | Rate of pH threshold shift | 0.01 /hr | * |
| - | Minimum pH threshold | 6.0 | [4] |
| - | Rate of proliferation probability shift | 0.001 /hr | * |
| - | Cell cycle duration | 30 hrs | [5] |
| $\boldsymbol{\lambda_{tolerance}}$ | Rate of acquiring tolerance to chemotherapeutic agent | 0.001 /hr | * |
| - | Duration of chemo exposure needed to induce tolerance | 0.02 cell cycle durations | [6] |
| - | Amount of chemo exposure needed to induce tolerance | 0.01 | [6] |
| $\boldsymbol{C_{tolerance}}$ | Initial amount of damage needed to kill a cell | 90 | * |
| $\boldsymbol{\lambda_C}$ | Chemotherapeutic agent uptake rate | 0.007 /s | * |
| $\boldsymbol{\lambda_{repair}}$ | Rate of damage repair | $1e^{-5}$ /s | * |
| - | Migration rate | [8.3, 24.9 (if hypoxic)] µm/s | [7] |
| - | Bias factor when selecting migration direction | 50 | * |

Vessel Parameters

| Symbol | Description | Value | Reference |
|---|---|---|---|
| - | Vessel diameter | 15 µm | [8,9] |
| $\boldsymbol{\mu}$ | Spring constant | 100 | * |

| | | | |
|---|---|---|---|
| - | Maximum overlap | 0.5 vessel diameters | * |
| - | Base vessel recruitment rate | [1.58e-6, 3.16e-6] /hr/μm$^2$ | * |
| $\boldsymbol{h_v}$ | Vessel health of recruited vessels | [0.25, 0.5] | * |

*Diffusion Parameters*

| Symbol | Description | Value | Reference |
|---|---|---|---|
| $\boldsymbol{V_O}$ | Maximum oxygen uptake rate (normal tissue) | 0.012 mmol/L/s | [3] |
| $\boldsymbol{k_O}$ | Half-max oxygen concentration (normal tissue) | 0.005 mmol/L | [3] |
| $\boldsymbol{k_G}$ | Half-max glucose concentration (normal tissue) | 0.04 mmol/L | [3] |
| $\boldsymbol{p_g}$ | Glycolytic shift (normal tissue) | 1.0 | [3] |
| $\boldsymbol{k_H}$ | Proton buffering coefficient (normal tissue) | 0.5e-4 | * |
| $\boldsymbol{C_{blood}}$ | Chemotherapeutic agent concentration in blood | 1 | [6] |
| $\boldsymbol{\lambda_{d,c}}$ | Chemotherapeutic decay rate | 0.005 /s | * |
| $\boldsymbol{D_C}$ | Chemotherapeutic diffusion coefficient | 50 μm$^2$/s | * |
| $\boldsymbol{D_O}$ | Oxygen diffusion coefficient | 1820 μm$^2$/s | [3] |
| $\boldsymbol{D_G}$ | Glucose diffusion coefficient | 500 μm$^2$/s | [3] |
| $\boldsymbol{D_H}$ | Proton (acid) diffusion coefficient | 1080 μm$^2$/s | [3] |
| $\boldsymbol{D_{VEGF}}$ | VEGF diffusion coefficient | 9.10 μm$^2$/s | [10] |
| $\boldsymbol{\lambda_p}$ | VEGF production | 1 /s | [10] |
| $\boldsymbol{\lambda_{d,V}}$ | VEGF degredation | 0.006 /s | [10] |
| $\boldsymbol{\lambda_{ves}}$ | VEGF removal by vasculature | 0.001 /s | [10] |
| - | Blood oxygen concentration | 0.056 mmol/L | [3] |
| - | Blood glycose concentration | 5.0 mmol/L | [3] |
| - | Blood pH | 7.4 | [3] |

* model specific, based on biological behavior

**REFERENCES**


[1] S.-J. Hao, Y. Wan, Y.-Q. Xia, X. Zou, and S.-Y. Zheng, *Size-based separation methods of circulating tumor cells*, Advanced Drug Delivery Reviews. 125 (2018) 3–20.

[2] J.M. Osborne, A.G. Fletcher, J.M. Pitt-Francis, P.K. Maini, and D.J. Gavaghan, *Comparing individual-based approaches to modelling the self-organization of multicellular tissues*, PLoS Computational Biology. 13 (2017) e1005387.

[3] M. Robertson-Tessi, R.J. Gillies, R.A. Gatenby, and A.R. Anderson, *Impact of metabolic heterogeneity on tumor growth, invasion, and treatment outcomes*, Cancer Research. 75 (2015) 1567–1579.

[4] M. Shamsi, M. Saghafian, M. Dejam, and A. Sanati-Nezhad, *Mathematical modeling of the function of Warburg effect in tumor microenvironment*, Scientific Reports. 8 (2018) 1–13.

[5] C.G. Cess and S.D. Finley, *Multi-scale modeling of macrophage—T cell interactions within the tumor microenvironment*, PLoS Computational Biology. 16 (2020) e1008519.

[6] J. Pérez-Velázquez and K.A. Rejniak, *Drug-induced resistance in micrometastases: analysis of spatio-temporal cell lineages*, Frontiers in Physiology. 11 (2020) 319.

[7] K.-A. Norton, K. Jin, and A.S. Popel, *Modeling triple-negative breast cancer heterogeneity: Effects of stromal macrophages, fibroblasts and tumor vasculature*, Journal of Theoretical Biology. 452 (2018) 56–68.

[8] J.C. Forster, W.M. Harriss-Phillips, M.J. Douglass, and E. Bezak, *A review of the development of tumor vasculature and its effects on the tumor microenvironment*, Hypoxia. 5 (2017) 21.

[9] J.R. Less, T.C. Skalak, E.M. Sevick, and R.K. Jain, *Microvascular architecture in a mammary carcinoma: branching patterns and vessel dimensions*, Cancer Research. 51 (1991) 265–273.

[10] G. Mahlbacher, L.T. Curtis, J. Lowengrub, and H.B. Frieboes, *Mathematical modeling of tumor-associated macrophage interactions with the cancer microenvironment*, Journal for Immunotherapy of Cancer. 6 (2018) 1–17.
